## Supplemental Information for "Cetacean coronavirus spikes highlight S glycoprotein structural plasticity"

*Cetacean coronavirus spikes highlight S glycoprotein structural plasticity* – **Supplementary Information**

### Table of Contents

**Supplementary Figure 1** Purification of BwCoV and BdCoV SED-GCN4-ST (trunc3) 2P proteins and negative stain data.

**Supplementary Figure 2** Cryo-EM data processing pipelines for BwCoV and BdCoV S.

**Supplementary Figure 3** Cryo-EM data processing of the BwCoV and BdCoV spike ectodomains

**Supplementary Figure 4** CeCoV S quaternary packing is reminiscent of alphaCoV spike proteins.

**Supplementary Figure 5** FoldTree analysis of coronavirus $S1^B$ domains supports ancestral relation of CeCoV $S1^B$ to alpha- and deltacoronaviruses.

**Supplementary Figure 6** AlphaFold 3 structure prediction models of the BdCoV and BwCoV $S1^0$ domains does not reflect experimentally determined structures.

**Supplementary Figure 7** CeCoV S displays distinctive elements in conserved S2 fusion machinery.

**Supplementary Figure 8** Detection of N-glycans on the surface of CeCoV spike glycoprotein.

**Supplementary Figure 9** Detection of O-glycans on the surface of the CeCoV spike glycoprotein.

**Supplementary Figure 10** Overview of experimentally detected N- and O-linked glycosylations of both CeCoV S proteins.

**Supplementary Figure 11** GammaCoV receptor-binding domain displays signs of recombination.

**Supplementary Table 1** Cryo-EM data collection, refinement and validation statistics for global refinements.

**Supplementary Table 2** Overview of unresolved regions in CeCoV N-terminal domains

**Supplementary Table 3** Overview of experimentally detected N-glycan abundance on coronavirus spike proteins

**Supplementary Table 4** Overview and conservation of predicted N-linked glycosites across CeCoV BwCoV and BdCoV S proteins along with the corresponding experimental data.

**Supplementary Data 1** FoldSeek_hits_BwCoV_S10.gz

**Supplementary Data 2** FoldSeek_hits_BdCoV_S10.gz

**Supplementary Data 3** BwCoV_AUC_per_glycosite_all_proteases.xlsx

**Supplementary Data 4** BdCoV_AUC_per_glycosite_all_proteases.xlsx

**Supplementary Data 5** N_glycol_site_detection_per_protease.xlsx

**Supplementary Data 6** O_glycans_Opair_pooledData.xlsx

**References**

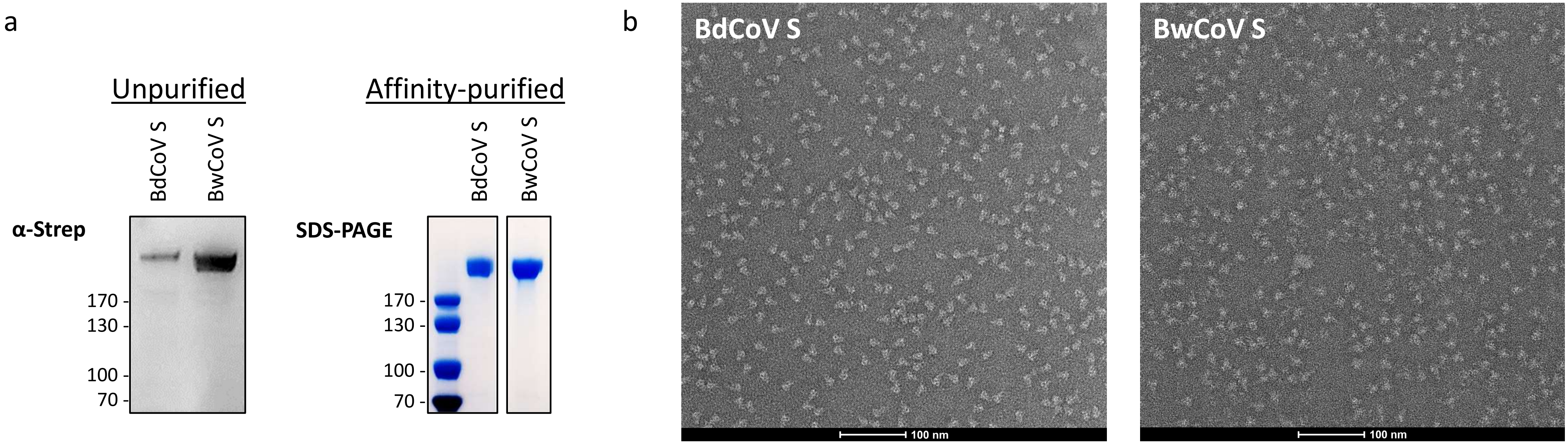

**Supplementary Figure 1** Purification of BwCoV and BdCoV SED-GCN4-ST 2P proteins and corresponding negative stain data.

a) Anti-strep Western blot and SDS-PAGE of CeCoV spike protein ectodomains used for cryoEM.

b) Negative stain micrographs of BwCoV and BdCoV spike ectodomains used for structural characterization. Micrographs display monodisperse particles that are of the expected size (~10 x 20 nm). Size bar represents 100 nm.

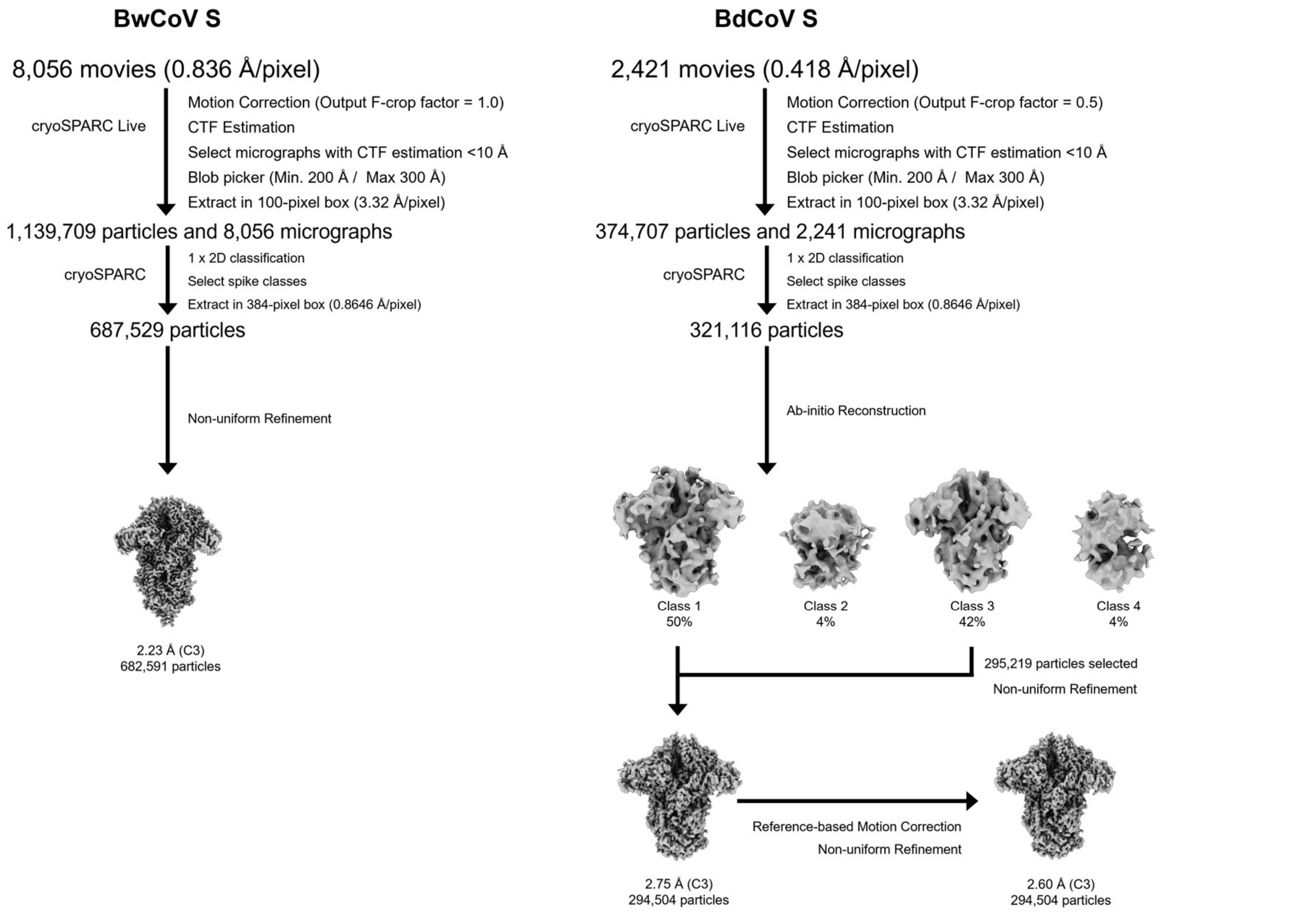

**Supplementary Figure 2** Cryo-EM data processing pipelines in cryoSPARC[10] for BwCoV (*left panel*) and BdCoV S (*right panel*).

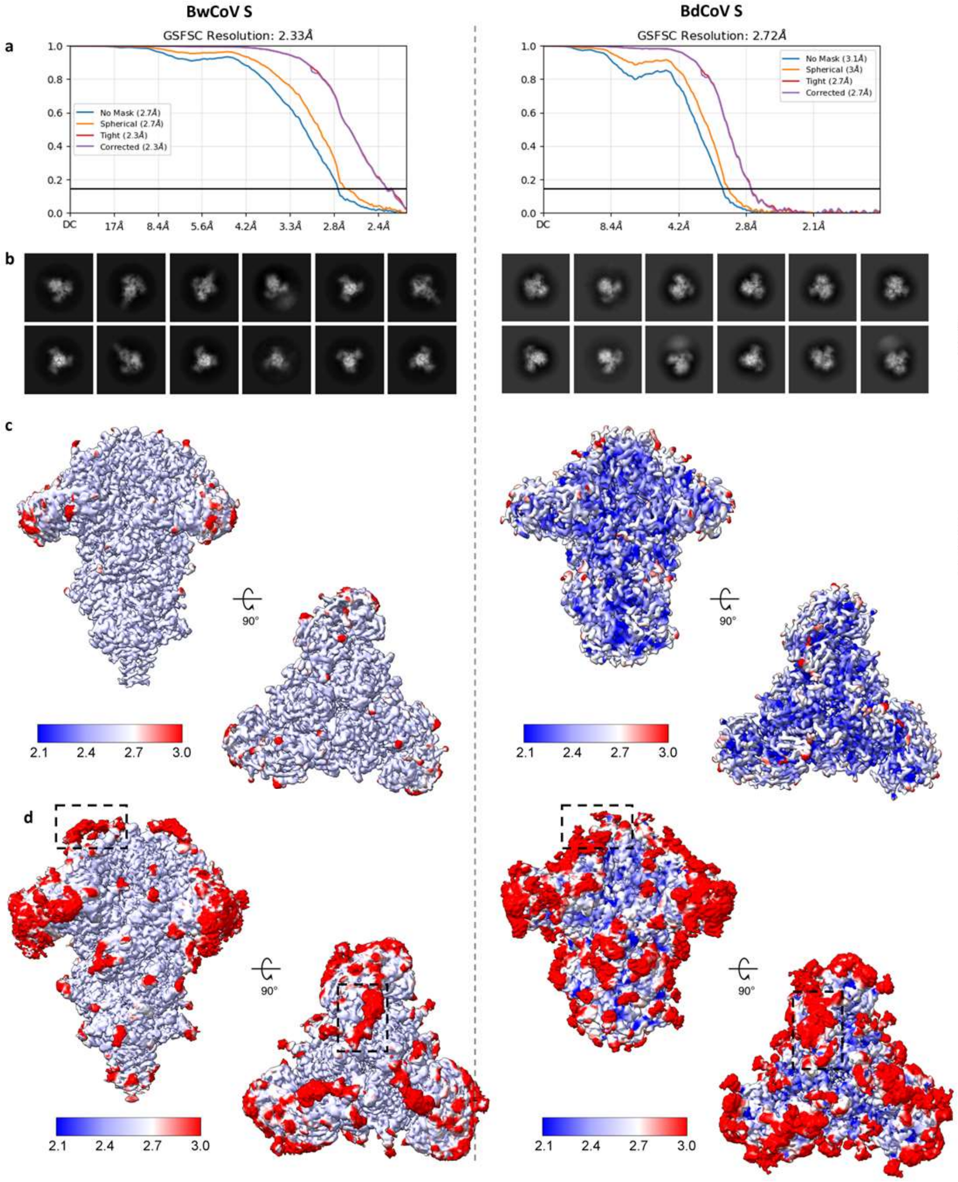

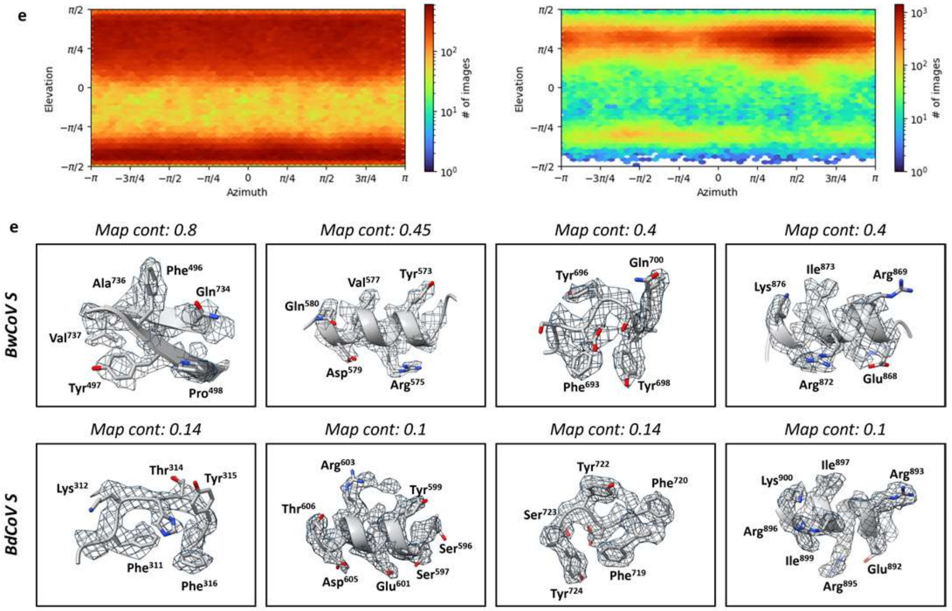

**Supplementary Figure 3** Cryo-EM data processing of the BwCoV and BdCoV spike ectodomains. a) FSC plots of BwCoV and BdCoV spike ectodomain reconstructions. Representative reference-free 2D class averages for BwCoV S. b) Tightly contoured EM density maps for the spike ectodomains coloured according to local resolution which was calculated in cryoSPARC[10] (countour level BwCoV: 0.156; BdCoV: 0.0544). c) Loosely contoured EM density maps for the spike ectodomains coloured according to local resolution which was calculated in cryoSPARC[10] (countour level BwCoV: 0.0931; BdCoV: 0.0206). Part of the flexible $S1^A$ loop decorated with O-glycans at the apex of the trimer can be observed at this contour level (dashed black boxes). d) Angular distribution plot calculated in cryoSPARC[10] for particle projections in the globally refined map. e) Examples of EM density + model to illustrate resolution of maps. Panels left to right: $S1^A$, $S1^B$, $S1^C$, S2.

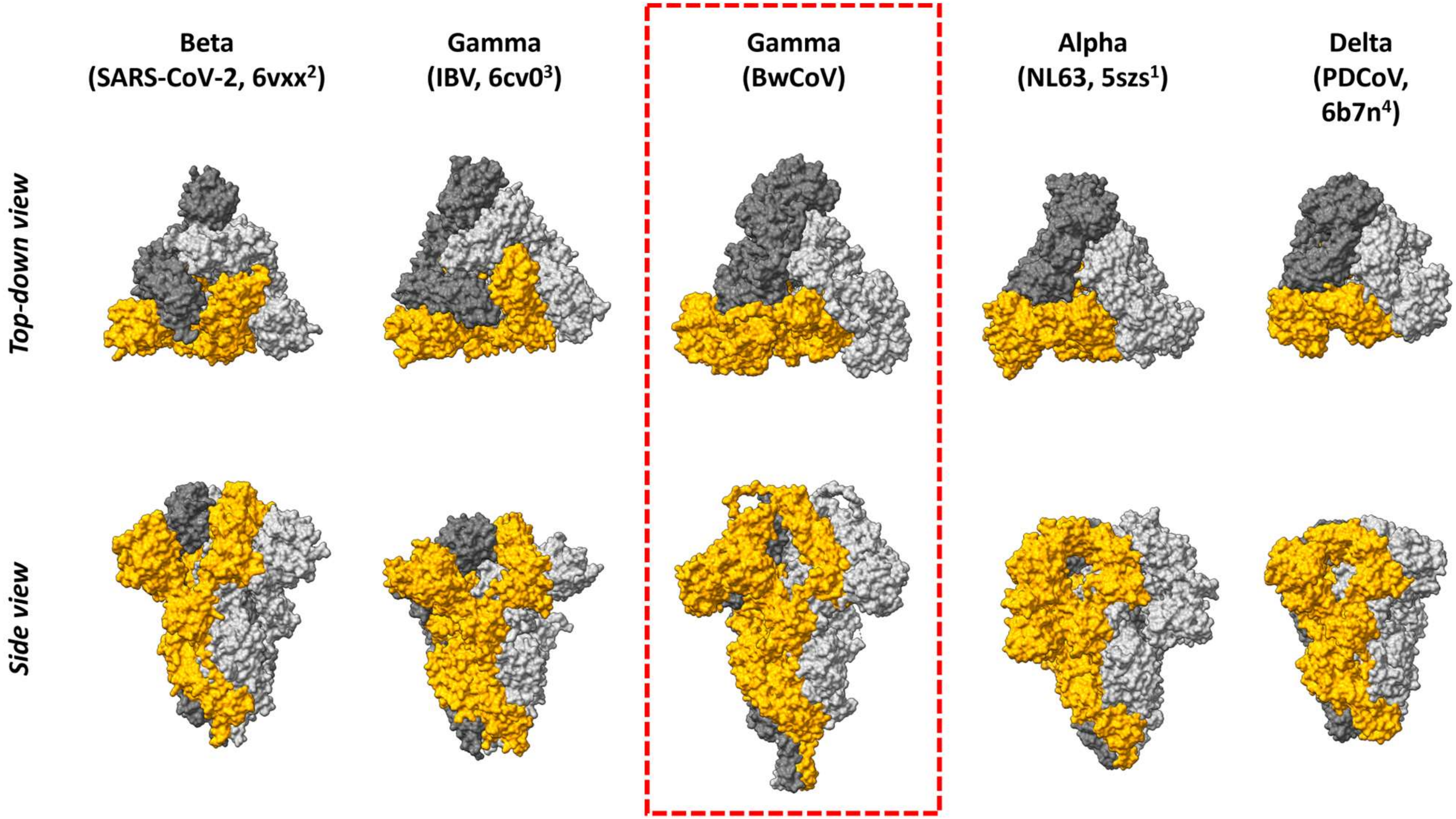

**Supplementary Figure 4** CeCoV S quaternary packing is reminiscent of alphaCoV spike proteins. Surface representations are shown of representative spike proteins from all four genera (IBV (6cv0), PDCoV (6b7n), SARS-CoV-2 (6vxx), HCoV-NL63 (5szs). Protomers of the trimeric models are coloured in the same manner for every protein. Models are ordered according to their packing mode: spike proteins displaying primarily inter-protomer contacts (beta-, gammaCoV) on the left and spike proteins displaying primarily intra-protomer contacts (alpha-, deltaCoV) on the right. The BwCoV spike structure is highlighted by a dashed red box.

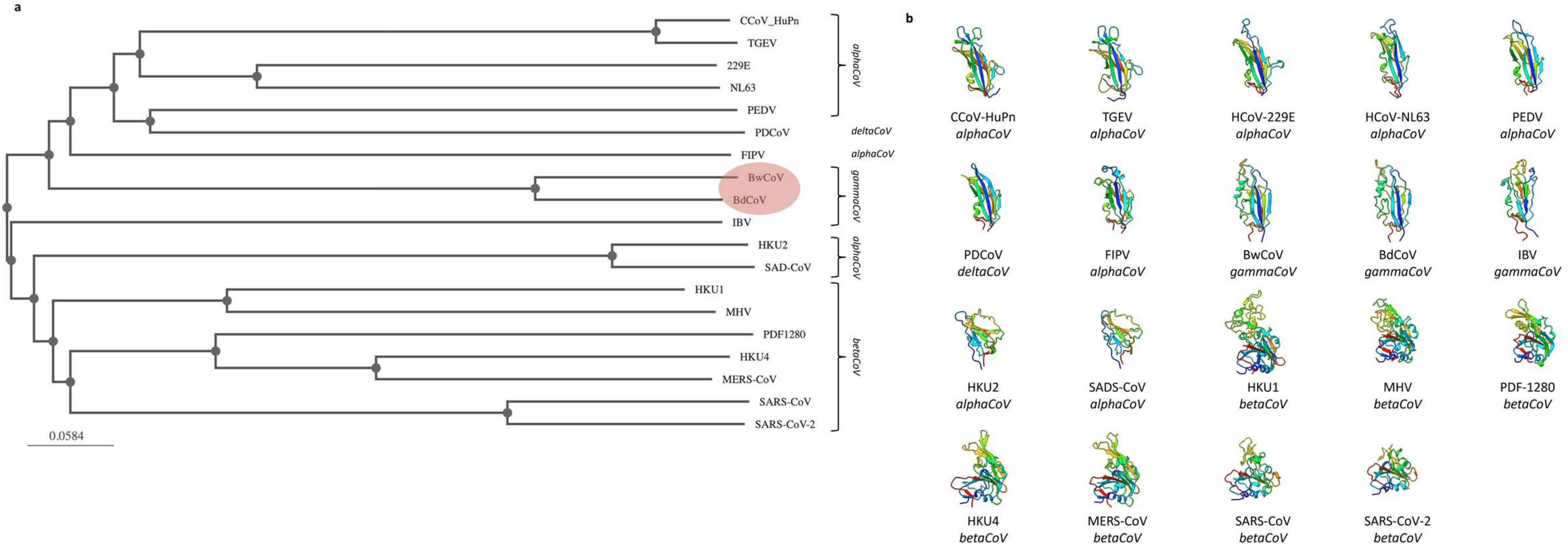

**Supplementary Figure 5** FoldTree[11] analysis of coronavirus S1$^{B}$ domains supports ancestral relation of CeCoV S1$^{B}$ to alpha- and deltacoronaviruses. A) FoldTree analysis of experimentally characterized coronavirus S1$^{B}$ domains. B) Cartoon representation of aligned and rainbow-coloured atomic models of experimentally characterized coronavirus S1$^{B}$ domains. The models are shown in the order of the phylogeny for comparison.

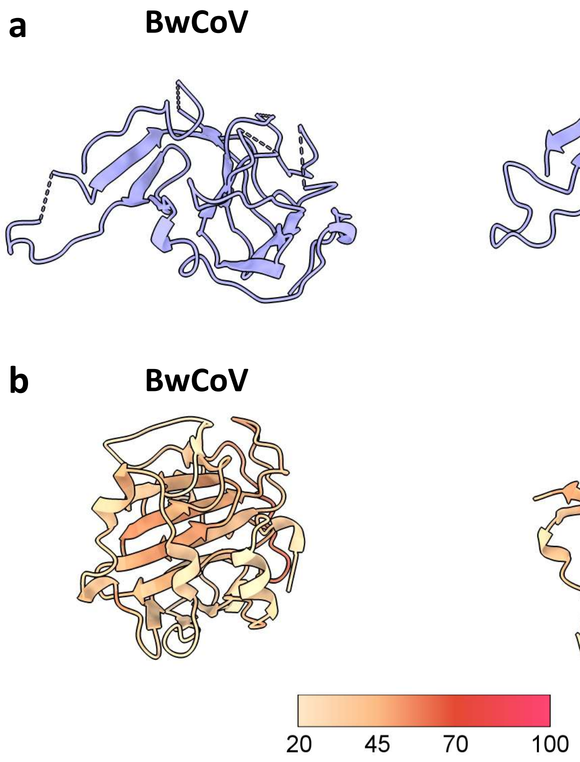

**Supplementary Figure 6. AlphaFold 3 structure prediction models of the BdCoV and BwCoV S1$^0$ domains does not reflect experimentally determined structures.** a) Cartoon representation of the experimentally determined structures of BdCoV and BwCoV S1$^0$, shown in the same (aligned) view. Unresolved regions are indicated by dashed lines, no glycans are shown for clarity. b) Cartoon representation of the top AlphaFold 3[14] models predicted for the amino acid sequences of domain S1$^0$ of the BdCoV and BwCoV spike proteins. Models were aligned to the experimentally determined structures shown in panel a and are coloured by pIDDT scores. The colour legend indicates the pIDDT confidence level (a higher score signifies a higher confidence).

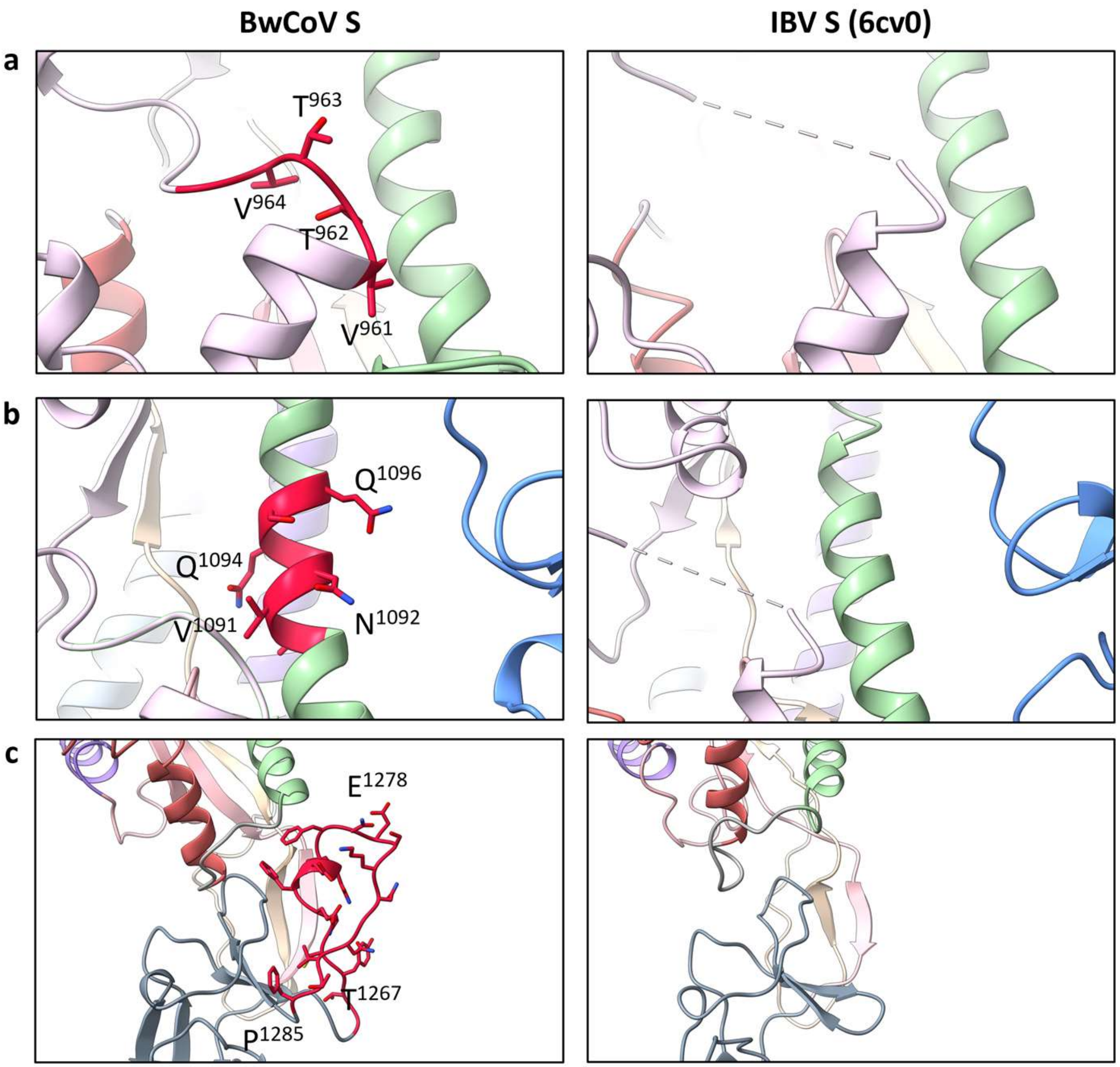

**Supplementary Figure 7 CeCoV S displays distinctive elements in conserved S2 fusion machinery.** a) Cartoon representation of the S2 stalk region of BwCoV (*left panel*) and IBV S (*right panel*) with the different S2 elements coloured as in Fig 1c for reference. The 5-residue insertion directly upstream of S2' site found in CeCoV S proteins (BwCoV residues Ser961 – Asp965; BdCoV residues Ser984 – Gly988) is shown in as red sticks coloured by heteroatom. b) The insertion of one heptad repeat in HR1 (BwCoV residues Thr1090 – Val1096; BdCoV residues Thr1113 – Val1119) is shown as red sticks coloured by heteroatom. c) The 21-residue insertion in HR2 (BwCoV residues Gly1266 – Phe1286; BdCoV residues Gly1289 – Phe1309) is shown as red sticks coloured by heteroatom.

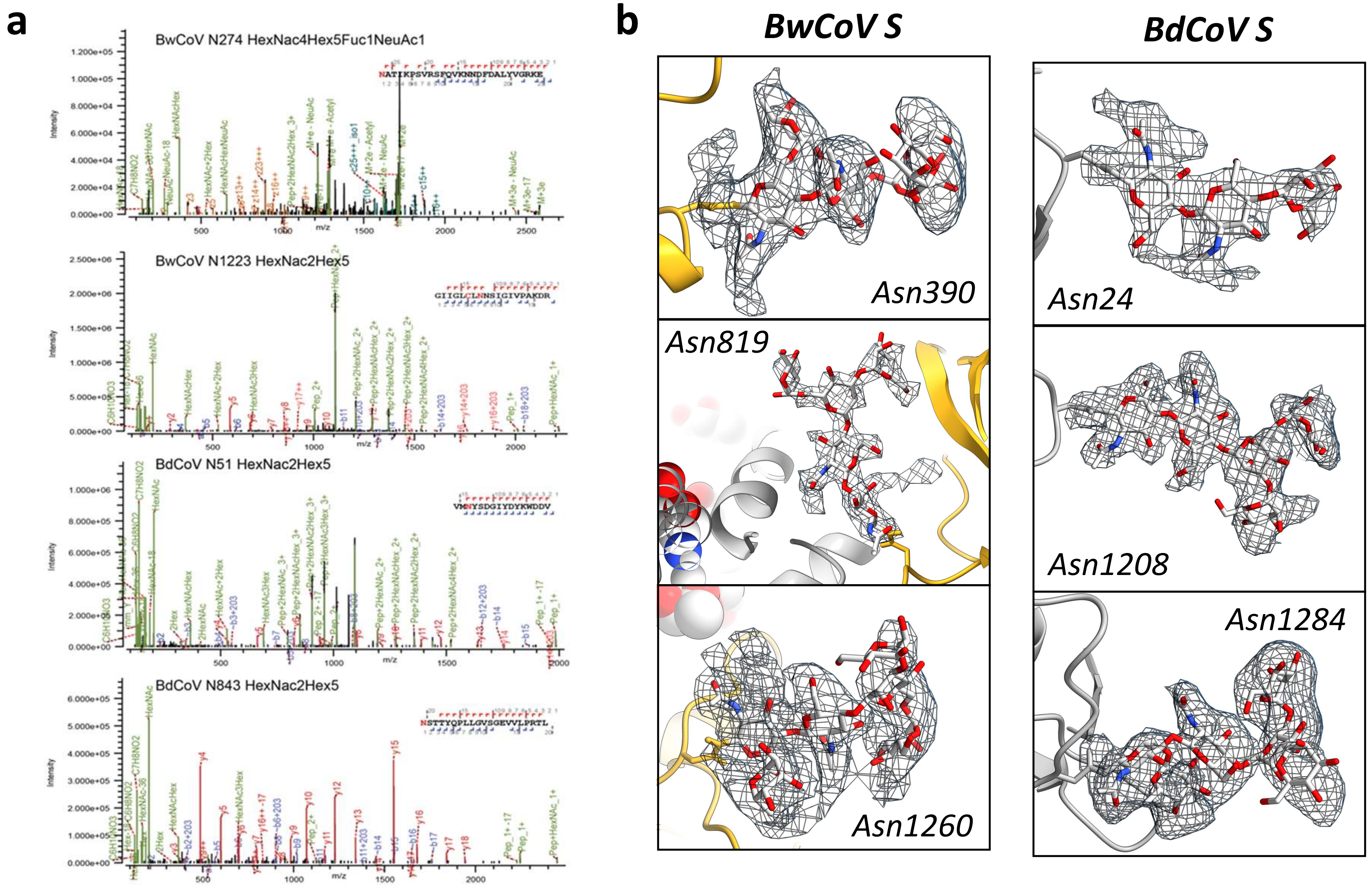

**Supplementary Figure 8 Detection of N-glycans on the surface of CeCoV spike glycoprotein.** a) Representative MS/MS spectra of the Byonic N-glycan identifications for BwCoV (top two panels) and BdCoV (lower two panels). b) EM density (grey mesh) zoned around the indicated modelled N-glycans of the BwCoV (*left panels*) and BdCoV (*right panels*) spike glycoproteins.

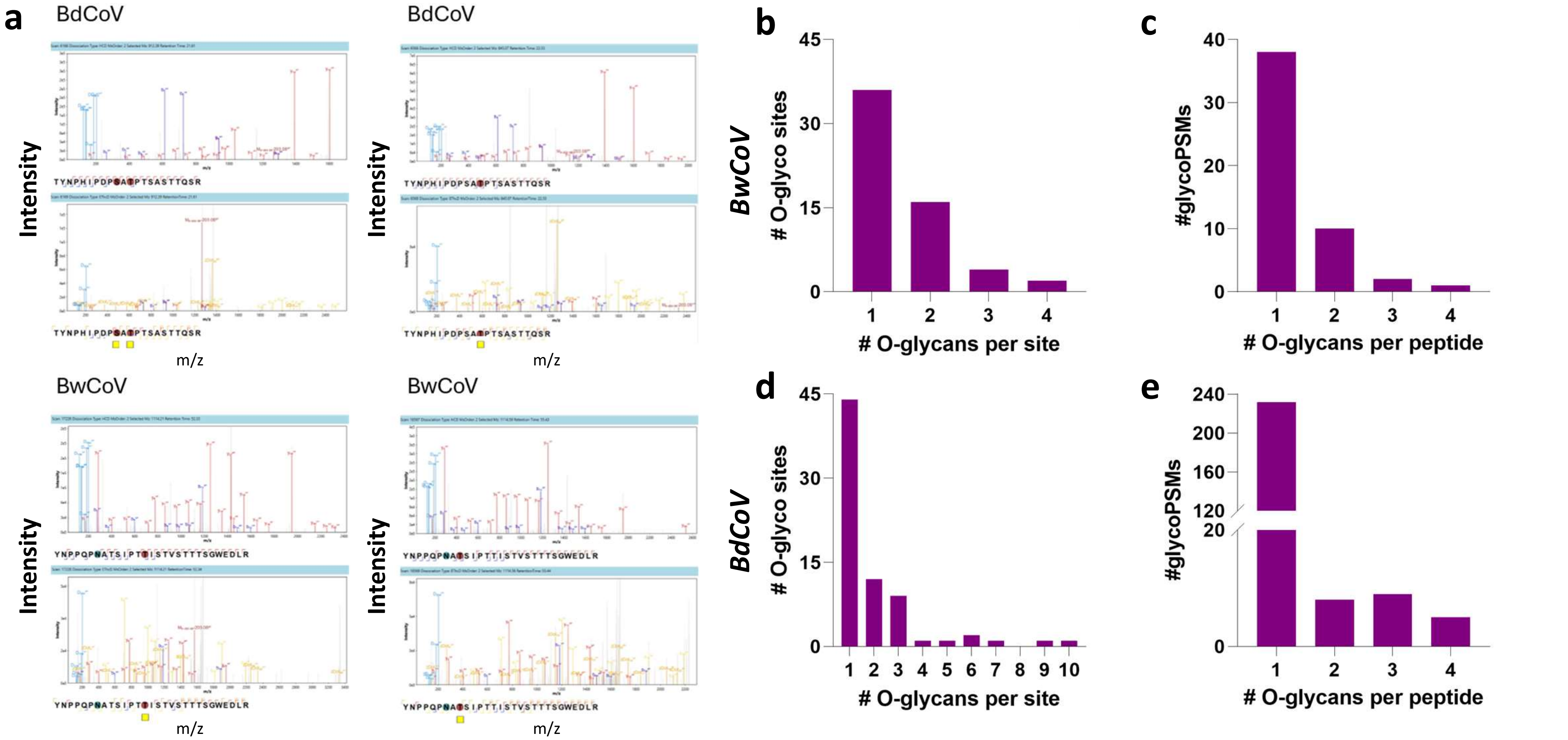

**Supplementary Figure 9 Detection of O-glycans on the surface of the CeCoV spike glycoprotein.** a) Representative MS/MS spectra supporting Opair identifications of O-glycosylation sites. Top spectra show HCD fragmentation (b- and y-ions; blue and red, respectively) and identification of oxonium ions (light blue); bottom spectra show EThcD fragmentation where additionally c- and z-ions can be seen (yellow and orange, respectively). b) and d) Number of identified O-glycosites with the number of identified O-glycoforms per site for BwCoV and BdCoV S, respectively. Most identified O-glycosites carry only one glycoforms per site. c) and e) Number of O-glycans identified on the same peptide.

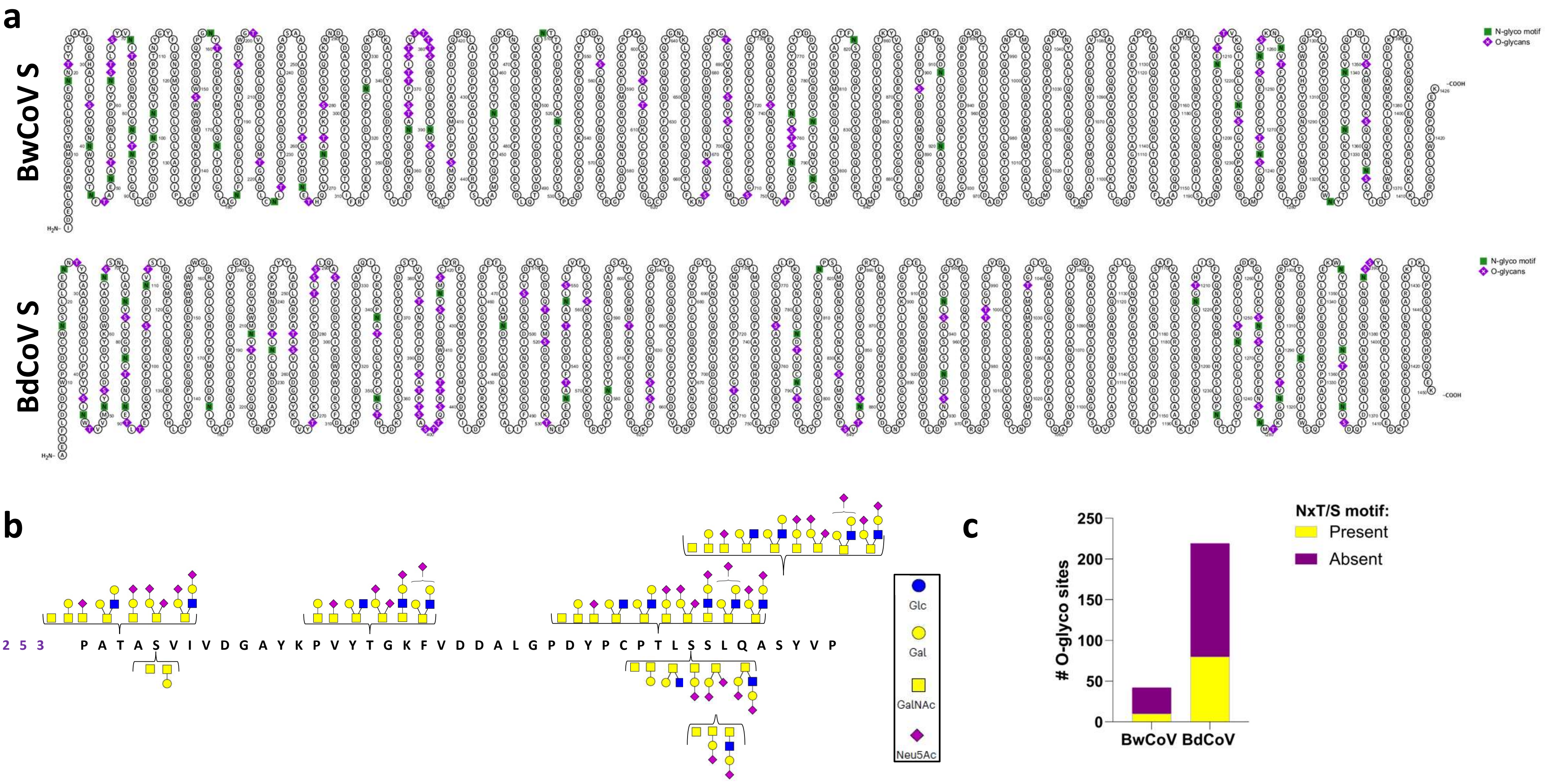

**Supplementary Figure 10 Additional O-glycosylation data.** a) Overview of experimentally detected N- and O-glycosylation patterns of both CeCoV S proteins. N-glycan sites are depicted in green and O-glycan sites in purple. b) Overview of dominant *O*-linked glycoforms within BdCoV residues 379-410 c) Number of O-glycosylation sites identified within an N-glycan motif for both CeCoV S proteins.

**Supplementary Table 1** Cryo-EM data collection, refinement and validation statistics for global refinements.

| | BwCoV S (EMDB-53512) (PDB 9R1Q) | BdCoV S (EMDB-53513) (PDB 9R1R) |
|---|---|---|
| **Data collection and processing** | | |
| Magnification | 105,000 x | 105,000x |
| Voltage (kV) | 300 | 300 |
| Electron exposure (e-/Å$^2$) | 50 | 50 |
| Defocus range (μm) | - | - |
| Pixel size (Å) | 0.83 | 0.83 |
| Symmetry imposed | C3 | C3 |
| Initial particle images (no.) | 1,139,709 | 374,707 |
| Final particle images (no.) | 682,591 | 294,504 |
| Map resolution (Å) | 2.2 | 2.6 |
| FSC threshold | 0.143 | 0.143 |
| Map resolution range (Å) | 2.18 - 20.69 | 1.81 - 15.95 |
| **Refinement** | | |
| Model resolution (Å) | | |
| FSC threshold 0.5 | 2.30 | 2.68 |
| FSC threshold 0.143 | 2.23 | 2.53 |
| Map sharpening *B* factor (Å$^2$) | 41.82 | 54.08 |
| **Model composition** | | |
| Non-hydrogen atoms | 38619 | 31422 |
| Protein residues | 3777 | 3663 |
| Ligands | 219 | 198 |
| **B factors (Å$^2$)** | | |
| Protein | 135.66 | 114.22 |
| Ligand | 130.61 | 119.10 |
| **R.m.s. deviations** | | |
| Bond lengths (Å) | 0.005 | 0.005 |
| Bond angles (°) | 0.75 | 0.69 |
| **Validation** | | |
| MolProbity score | 1.95 | 1.74 |
| Clashscore | 21.00 | 9.02 |
| Poor rotamers (%) | 0.00 | 0.28 |
| **Ramachandran plot** | | |
| Favoured (%) | 97.26 | 96.14 |
| Allowed (%) | 2.65 | 3.53 |
| Disallowed (%) | 0.08 | 0.33 |

**Supplementary Table 2** Overview of unresolved regions in CeCoV N-terminal domains. Topologically equivalent regions of the two spike proteins are listed within the same row.

| Domain | BwCoV spike | BdCoV spike |
|---|---|---|
| $S1^0$ | 57-63 | |
| | 85-96 | 91-101 |
| | 106-115 | 112-114 |
| | 151-162 | 160-165 |
| | 189-196 | 193-204 |
| $S1^A$ | | 226-230 |
| | 265-276 | 277-303 |
| | | 392-414 |
| S2 | 776-784 (around S1/S2 junction) | 801-812 (around S1/S2 junction) |
| | 1332-end | 1333-end |

**Supplementary Table 3** Overview of experimentally detected N-glycan abundance on coronavirus spike proteins. N-glycosylation calculated as 2.5 kDa addition to MW of glycoprotein.

| Glycoprotein | N-glycosylation Sequons (kDa) | # protein residues (kDa) | N-glycan contribution to total MW | Reference |
|---|---|---|---|---|
| **HIV-1 gp** | 31 (77.5) | 859 (95) | 44.9% | Kwon, 2015, *NSMB*[5] |
| **BwCoV S** | 37 (92.5) | 1449 (160) | 36.6% | *This study* |
| **HCoV-NL63 S** | 34 (85) | 1356 (149) | 36.2% | Walls, 2016, *NSMB*[1] |
| **BdCoV S** | 35 (87.5) | 1472 (162) | 35.4% | *This study* |
| **LASV gp** | 11 (27.5) | 491 (54) | 33.7% | Hastie, 2017, *Science*[6] |
| **HCoV-HKU1 S** | 29 (72.5) | 1356 (149) | 32.6% | Watanabe, 2020, *Nat Comm*[7] |
| **SARS-CoV S** | 23 (57.5) | 1255 (138) | 29.1% | Watanabe, 2020, *Nat Comm*[7] |
| **SARS-CoV-2 S** | 22 (55) | 1273 (140) | 28.2% | Watanabe, 2020, *Science*[8] |
| **MERS-CoV** | 23 (57.5) | 1353 (149) | 27.7% | Watanabe, 2020, *Nat Comm*[7] |
| **IAV HA (H1)** | 9 (22.5) | 566 (62) | 26.6% | Lee, 2014, *Nat Comm*[9] |

| | BwCoV S | | | | BdCoV S | | | |
|---|---|---|---|---|---|---|---|---|
| | *Sequon* | *Glycan detected by MS?* | *Protein region resolved in EM?* | *EM density for glycan chain?* | *Sequon* | *Glycan detected by MS?* | *Protein region resolved in EM?* | *EM density for glycan chain?* |
| **1** | | | | | 17 – NSS | complex | Yes | Yes |
| **2** | 19 – NNT | mannose | Yes | Yes | 24 – NNT | mannose | Yes | Yes |
| **3** | 40 – NWT | mannose | Yes | Yes | 45 – NWT | mannose | Yes | Yes |
| **4** | 46 – NFT | mannose | Yes | Yes | 51 – NYS | mannose | Yes | Yes |
| | 51 – NAT | complex | Yes | No | | | | |
| **5** | 63 – NST | complex | No | No | 67 – NSS | complex | Yes | No (only low res) |
| **6** | 71 – NIT | mannose | Yes | Yes | 75 – NVS | mannose | Yes | Yes |
| **7** | 82 – NFT | mannose | Yes | Yes | 82 – NCT | mannose | Yes | Yes |
| **8** | 85 – NRT | ~none | No | No | 88 – NET | complex | Yes | Yes |
| | 100 – NCT | mannose | Yes | Yes | | | | |
| **9** | 103 – NVT | ~none | Yes | No | 110 – NVT | complex | Yes | No (only low res) |
| | 158 – NYT | complex | No | No | | | | |
| **10** | 172 – NIT | complex | Yes | Yes | 176 – NIT | mannose | Yes | Yes |
| | 181 – NFS | mannose | Yes | Yes | | | | |
| **11** | | | | | 211 – NVT | mannose | Yes | Yes |
| **12** | 224 – NLT | complex | Yes | Yes | 234 – NLT | complex | Yes | Yes |
| | 265 – NET | complex | No | No | | | | |
| | 274 – NAT | complex | No | No | | | | |
| **13** | | | | | 341 – NAT | mannose | Yes | Yes |
| **14** | 326 – NET | complex | Yes | Yes | 351 – NET | complex | Yes | Yes |
| | 365 – NAT | mannose | Yes | Yes | | | | |
| **15** | 390 – NMS | complex | Yes | Yes | 417 – NMS | complex | Yes | Yes |
| **16** | | | | | 474 – NDT | mannose | Yes | Yes |
| | 455 – NYT | mannose | Yes | Yes | | | | |
| **17** | 509 – NTT | complex | Yes | Yes | 536 – NAT | complex | Yes | Yes |
| **18** | 521 – NLT | hybrid | Yes | Yes | 548 – NLS | mannose | Yes | Yes |
| **19** | | | | | 581 – NYT | complex | Yes | Yes |
| **20** | 757 – NAS | complex | Yes | Yes | 783 – NDT | mannose | Yes | Yes |
| **21** | 763 – NTT | complex/mannose | Yes | Yes | 789 – NIT | complex | Yes | Yes |
| | 785 – NVT | hybrid | Yes | No | | | | |
| **22** | 819 – NST | mannose | Yes | Yes | 843 – NST | mannose/hybrid | Yes | Yes |
| **23** | 911 – NIS | hybrid | Yes | Yes | 935 – NIS | complex | Yes | Yes |
| **24** | 920 – NAS | complex | Yes | Yes | 944 – NDS | complex | Yes | Yes |
| **25** | | | | | 1208 – NGT | mannose | Yes | Yes |
| **26** | 1209 – NET | mannose | Yes | Yes | 1233 – NET | mannose | Yes | Yes |
| **27** | 1223 – NNS | complex | Yes | Yes | 1247 – NNS | mannose | Yes | Yes |
| **28** | 1242 – NGT | complex | Yes | Yes | 1266 – NGS | mannose | Yes | Yes |
| **29** | 1254 – NES | mannose | Yes | Yes | 1278 – NMT | mannose | Yes | Yes |
| **30** | 1260 – NVT | complex | Yes | Yes | 1284 – NVT | complex | Yes | Yes |
| **31** | | | | | 1314 – NSS | mannose | Yes | Yes |
| **32** | 1323 – NYT | complex | Yes | Yes | 1347 – NYT | ~none | No | no |
| **33** | 1333 – NVT | none | No | No | 1357 – NVT | mannose | No | no |
| **34** | 1340 – NVS | none | No | No | 1364 – NVS | ~none | No | no |
| **35** | 1363 – NSS | complex | No | No | 1387 – NSS | ~none | No | no |

**Supplementary Table 4** Overview and conservation of predicted N-linked glycosites across CeCoV BdCoV and BwCoV S proteins along with their corresponding experimental data.

*Cetacean coronavirus spikes highlight S glycoprotein structural plasticity* – **Supplementary Information**
